# Nonsense, but not frameshift, truncating mutations result in Cyclin D2 stabilisation in an induced pluripotent stem cell model of MPPH

**DOI:** 10.1101/2025.08.29.673076

**Authors:** Erica L. Harris, Rowan D. Taylor, Katarzyna Szymanska, Ailsa M.S. Rose, Jacquelyn Bond, Colin A. Johnson, James A. Poulter

## Abstract

Stabilisation of Cyclin D2 is the underlying cause of a range of neurodevelopmental disorders, characterised by megalencephaly, cortical migration defects and overgrowth. Intracellular CCND2 is vital for mTOR pathway signalling, with inhibition of mTOR resulting in CCND2 phosphorylation and degradation by the ubiquitin proteasome system. Mutations at the regulatory c-terminus of CCND2, and in proteins that regulate mTOR such as PTEN, PIK3CA, AKT3 and TSC1/2, result in CCND2-stabilisation and overgrowth. To determine the molecular and cellular mechanisms underpinning the neurodevelopmental defects observed in Megalencephaly-Polymicrogyria-Polydactyly-Hydrocephalus (MPPH) syndrome, we generated human induced pluripotent stem cell (iPSC) derived models of CCND2-associated disease. Using CRISPR-Cas9 we generated lines containing either a pathogenic CCND2 variant (c.814G>T, p.Glu272Ter) or frameshift variants in the final exon of *CCND2*, all of which truncate CCND2 before the critical Thr-280 residue, required for its phosphorylation and degradation. We observed truncating frameshift variants do not result in CCND2 stabilization, whereas the single nucleotide c.814G>T, p.Glu272Ter substitution does, mimicking the effect seen in MPPH patients. Differentiation into human cortical spheroids (hCS) revealed all CCND2-truncating lines continued to express PAX6 beyond the neural progenitor (NP) expansion phase. Furthermore, both the homozygous and heterozygous p.Glu272Ter hCS failed to produce mature Tbr-1 expressing neurons, while some expression was observed in the frameshift hCS, highlighting differences in neurogenesis between frameshift and nonsense lines. Despite all lines truncating CCND2 and removing Thr-280, our data implies that frameshift truncations do not stabilise CCND2. In comparison, truncation of CCND2 through introduction of a single nucleotide nonsense variant results in CCND2 stabilisation, mimicking MPPH.

## Introduction

Mutations in genes encoding components of the phosphatidylinositol-3-kinase (PI3K)-AKT pathway, result in a genetically and phenotypically heterogenous group of brain overgrowth and malformation disorders (Riviere et al., 2012, Mirzaa et al., 2014, Lee et al., 2012, Lindhurst et al., 2012, Poduri et al., 2012, Mirzaa et al., 2012, Clayton-Smith et al., 1997, Mirzaa et al., 2004, Conway et al., 2007). Germline *de novo* mutations in *CCND2* cause megalencephaly-polymicrogyria-polydactyly-hydrocephalus (MPPH) syndrome, caused by substitution or deletion of residues at the c-terminus of CCND2, in particular Thr-280 and Pro-281, resulting in CCND2 stabilisation (Mirzaa et al., 2014). As all MPPH mutations identified to date are in the final exon, all premature termination mutations are predicted to result in a truncated protein that escapes nonsense mediated decay (Lindeboom et al., 2019) and lacks the key regulatory residues, specifically, Thr-280. Thus, Cyclin D2 accumulates resulting in a failure to exit the cell cycle at the G1/S checkpoint, leading to continued cellular proliferation. This is believed to be the underlying disease mechanism, with an accumulation of neural progenitor (NP) cells in the sub-ventricular zone of the developing neo-cortex, resulting in the clinical phenotypes observed in MPPH (Glickstein et al., 2009).

Differentiation of induced pluripotent stem cells (iPSCs) into three-dimensional brain organoids can reflect key human brain structures and development processes more accurately than 2D models. Human cortical spheroids (hCSs), for example, can mirror the human cerebral cortex and corticogenesis patterning seen in mid-fetal brain development (19-24 post conception weeks)(Pasca et al., 2015). This is achieved in exclusively nonadherent conditions, without extracellular scaffolding, through SMAD pathway inhibition and introduction of neurotrophic factors BDNF and NT3. With the ability to generate both deep and superficial layer-specific cortical neurons (Otani et al., 2016), as well as astrocytes (Blair et al., 2018), these 3D culture methods have been used to model a range of neurodevelopmental disorders (Blair et al., 2018, Xu et al., 2019, Birey et al., 2022, Balogh et al., 2024).

Here we use CRISPR/Cas9 to introduce truncating Cyclin D2 mutations into induced pluripotent stem cells to model MPPH. Whereas knock-in of a pathogenic nonsense variant, p.Glu272Ter, mimics MPPH through CCND2 stabilisation, similar protein truncations introduced by frameshift mutations do not. IPSC lines containing CCND2 truncations showed altered corticogenesis, with an increase in PAX6^+^ expressing NP cells in day 60 cortical spheroids and reduced expression of cortical neuron markers, such as TBR-1. Despite all lines truncating CCND2 and removing Thr-280, our data implies that frameshift truncations do not stabilise CCND2. In comparison, truncation of CCND2 through introduction of a single nucleotide nonsense variant results in CCND2 stabilisation, mimicking MPPH in an iPSC-derived organoid model.

## Results

### Generation of truncated CCND2 crispant iPSCs

To generate CCND2-truncated human iPSCs, we first used CRISPR-Cas9 without a repair template to introduce frameshifts within the final exon of *CCND2.* We hypothesised this would result in a truncated protein lacking Threonine-280, that escapes nonsense mediated decay due to the lack of an exon-exon junction following the mutation. Following transfection of the px458 CRISPR/Cas9 plasmid, containing a sgRNA targeting *CCND2* exon 5, we identified an edited (crispant) colony with compound heterozygous frameshift deletions (Fig. 1A), herein referred to as Cyclin D2 frameshift truncation line 1 (CCND2^FS-T1^). TOPO cloning followed by Sanger sequencing identified a 23-bp deletion, resulting in a frameshift and premature termination codon at residue 266 (c.785_809del23, p.Q263Vfs*4) on one allele, and a 5-bp deletion on the 2nd allele, resulting in a frameshift and premature termination at residue 272 (c.797_804del5, p.D267Ifs*6) (Fig. 2B). These mutations were predicted to encode truncated proteins similar in size to known human disease-associated mutations (e.g. p.Lys270Ter)(Mirzaa et al., 2014) and lacking the Thr280 residue required for phosphorylation and ubiquitination-dependent degradation (Fig. 2A).

**Figure 1:**
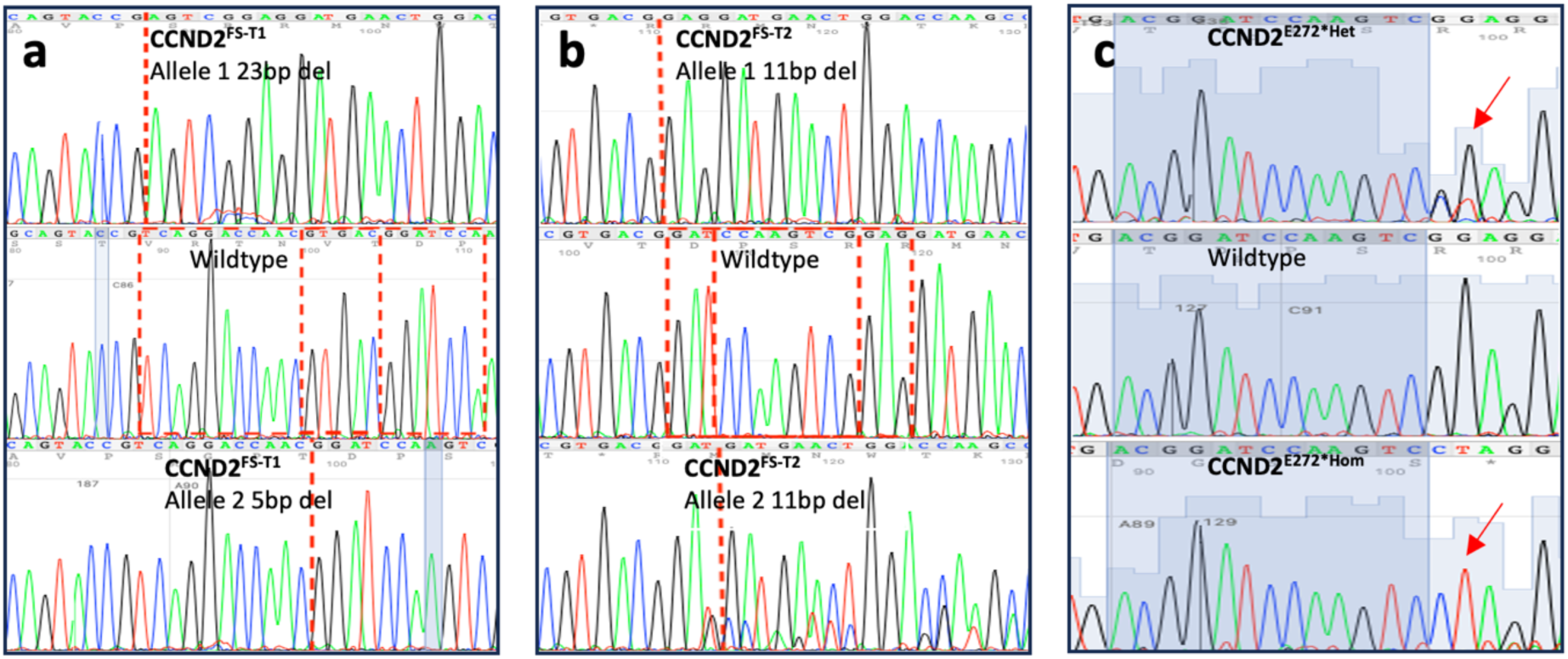
Creation of CRISPR-Cas9 engineered CCND2-truncating iPSC lines. A - Electropherogram showing both alleles (top and bottom) of CCND2 biallelic frameshift line CCND2^FS-T1^ compared to wildtype (middle) causing truncation at positions 266 and 272 on allele 1 and allele 2, respectively. B - illustrates the same but for the second CCND2 biallelic frameshift line CCND2^FS-T2^ showing an 11 bp deletion on allele 1 (top) and a different 11 bp deletion on allele 2 (bottom) compared to wildtype (middle), causing CCND2 truncation at position 272 and 274 respectively. C - Sanger sequencing electropherograms indicating with arrows the single base pair change c.814G>T in heterozygous (top) and homozygous (bottom) CCND2^E272*Het^ and CCND2^E272*Hom^ CRISPR-Cas9 edited lines compared to wildtype (middle).

**Figure 2.**
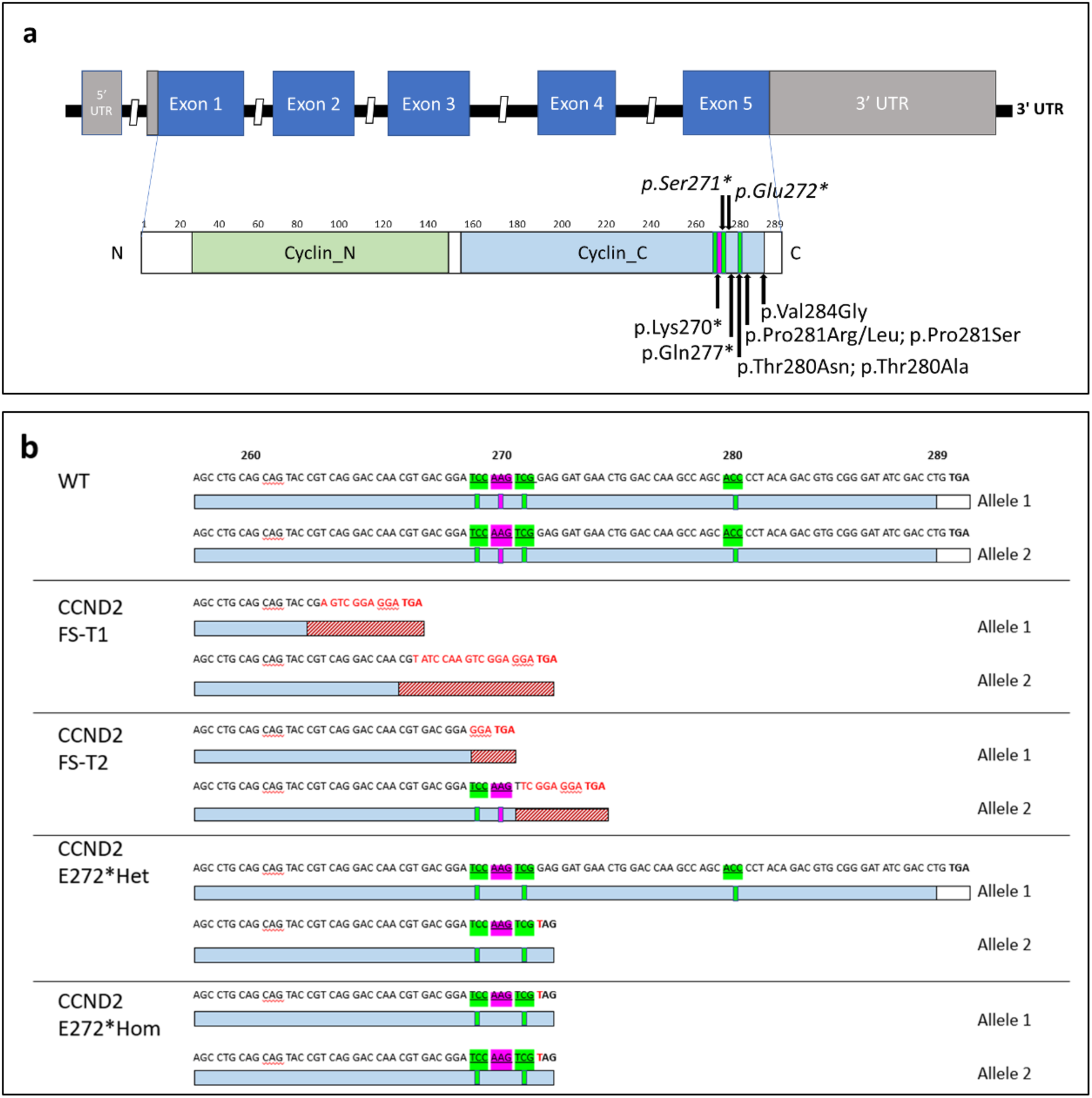
2D genomic and proteomic structure of CCND2 and CRISPR-Cas9 engineered truncation lines. A - 2D genomic and proteomic structure of CCND2, showing location of MPPH disease causing variants at the C-terminus (bottom) (Mirzaa *et al*., 2014, Zhao *et al*., 2024) and likely pathogenic variants from ClinVar in italics (top). Green indicates phosphorylation sites Ser269, Ser271 and Thr280, pink indicates ubiquitination site Lys270. B – Illustration of the CCND2 C- terminus showing the site of truncations, at the gene and protein level, in each CRISPR-Cas9 engineered line in comparison to wildtype (WT). Red hatched regions indicate regions of amino acid sequence changes due to a frameshift mutation. Green and pink highlighted regions signify locations of phosphorylation sites (Ser269, Ser271 and Thr280) and ubiquitination site Lys270, respectively.

A second crispant iPSC line, herein referred to as CCND2^FS-T2^, was created using the same method as CCND2^FS-T1^ however the esCas9 plasmid was used. This resulted in an iPSC line containing similar compound heterozygous frameshift mutations (Fig. 1B). Sanger sequencing after TOPO cloning confirmed these as being an 11 bp deletion (c.805_815del11, p.S269Gfs*1) on one allele and a different 11 bp deletion (c.811_821del11, p.S271Ffs*3) on the second allele, leading to truncation of CCND2 at positions 270 and 274 respectively. As with CCND2^FS-T1^, the mutations resulted in the translation of a truncated CCND2 protein lacking the regulatory Thr280 residue (Fig. 2B)

As a comparison to the frameshift lines, we also generated isogenic iPSC lines containing a pathogenic MPPH nonsense mutation: c.814G>T, p.Glu272Ter (ClinVar: VCV000985457.3). Using the same sgRNA used to create the two frameshift lines, we used a repair ssODN with the eCas9 plasmid to introduce the c.814G>T variant by homology directed repair, resulting in heterozygous and homozygous lines, herein referred to as CCDN2^E272*Het^ and CCND2^E272*Hom^ (Fig. 1C). Altogether we generated four models of CCND2 truncation, two frameshift lines (CCND2^FS-T1^ and CCND2^FS-T2^), and two SNV lines (CCND2^E272*Het^ and CCND2^E272*Hom^), all of which truncated CCND2 prior to the regulatory Thr-280 residue (Fig. 2B). Immunofluorescence staining of nuclear Oct3/4 and the cell surface marker SSEA was observed in all lines confirming they had retained their pluripotency and morphology (Fig. 3A).

**Figure 3.**
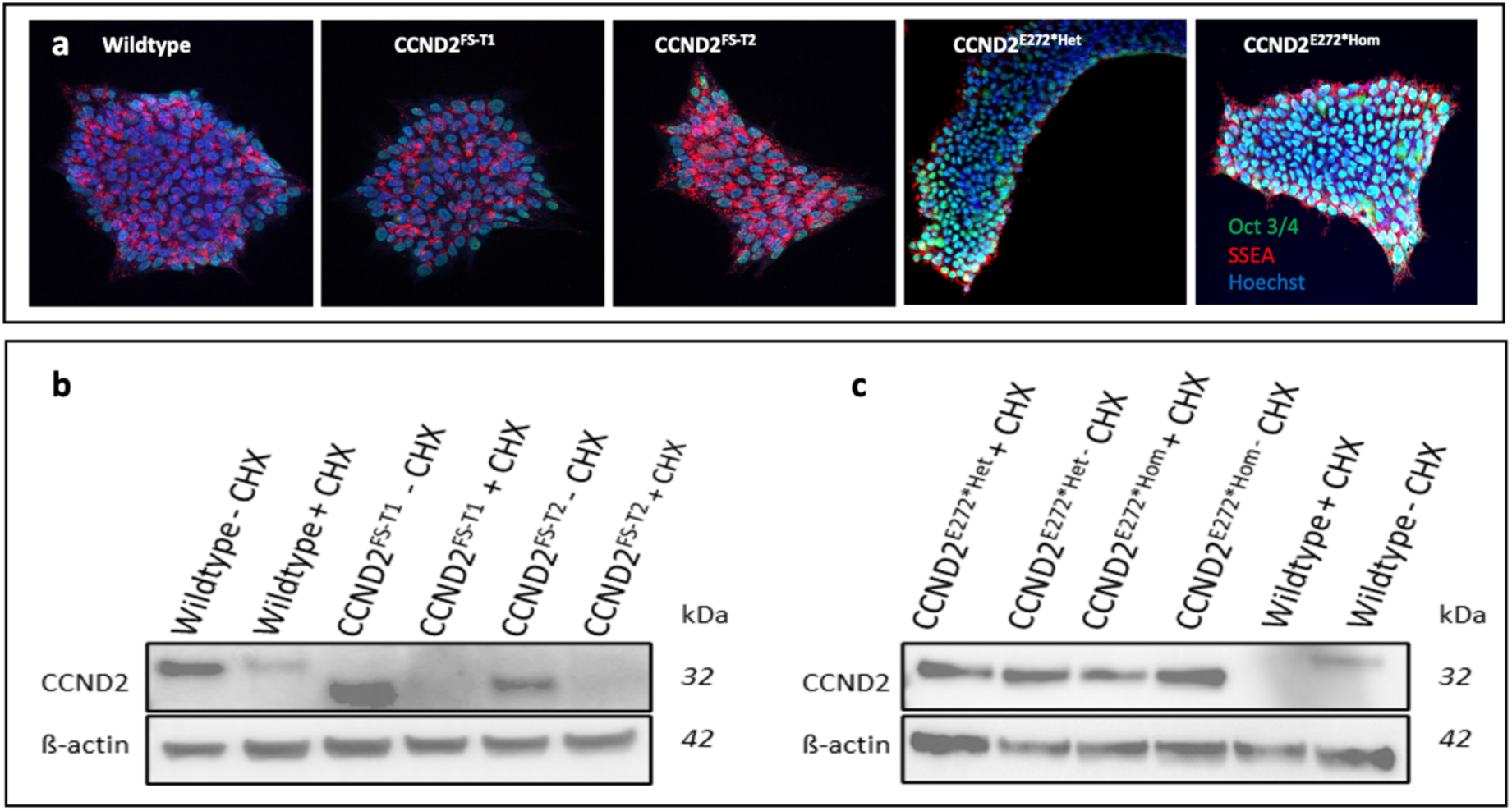
Induced pluripotent stem cell (iPSC) lines harbouring CCND2 truncation mutations express pluripotency markers, but only nonsense single nucleotide mutations cause CCND2 stabilisation. A - Representative fluorescence images of pluripotency markers on wildtype and mutant iPSC lines, stained for Oct 3/4 (green), SSEA4 (red) and Hoechst (blue). B - Representative western blot of CCND2 in the presence or absence of cycloheximide in CCND2^FS-T1^ and CCND2^FS-T2^ frameshift lines. C - Representative western blot of CCND2 in the presence or absence of cycloheximide in CCND2^E272*Het^ and CCND2^E272*Hom^ nonsense lines.

### Frameshift mutations do not cause CCND2 stabilisation

Previous studies have shown that MPPH causing mutations result in CCND2 stabilisation (Mirzaa et al., 2014). Because of the location of the variants we introduced, we hypothesized all four lines would express a truncated version of CCND2 without the regulatory Thr-280 residue (Fig. 2A) and therefore would result in CCND2 stabilisation. We therefore assessed CCND2 degradation following inhibition of protein translation with cycloheximide (CHX) in all cell lines and the parental unedited (wildtype) line. Unexpectedly, CCND2 stabilisation was not observed in either frameshift line, 90 minutes after addition of CHX (Fig. 3B). Furthermore, no increase in CCND2 levels were observed in either line compared to the wildtype, with the CCND2^FS-T2^ iPSC line showing significantly less CCND2. In both lines, protein truncation was confirmed (Fig. 3B).

In contrast, both the CCND2^E272*Het^ and CCND2^E272*Hom^ iPSCs had significantly higher levels of CCND2 compared to the wildtype and were confirmed to be truncated (Fig. 3C). In both lines, CCND2 stabilization was observed following treatment with cycloheximide, with no evidence of protein degradation compared to wildtype (Fig. 3C). Altogether, our data suggests that frameshift truncation mutations in the last exon of *CCND2* do not result in CCND2 stabilisation, compared to comparable truncations caused by null (single nucleotide) mutations.

### CCND2 mutations impair neurogenesis in human cortical spheroid models

To investigate how the different CCND2 truncating mutations affect neurodevelopment in a relevant 3D model system, we differentiated wildtype and crispant iPSCs into human cortical spheroids (hCS) using an established protocol (Pasca et al., 2015). We assessed the progress of cortical spheroid differentiation at two time points, day (D) 25 and D60, by immunofluorescent (IF) staining of cell-type specific markers for neural progenitor cells and neurons.

Staining of sectioned D25 spheroids confirmed the presence of PAX6 expressing NP cells in all CCND2 crispant lines, with little difference in expression observed compared to the wildtype (Fig. 4). Interestingly, both CCND2^E272*^ lines also had more ß-Tubulin III (TUJ1) expressing cells, suggestive of some premature neuronal differentiation. Following the neural progenitor expansion phase, PAX6 expression was reduced in wildtype spheroids, which coincided with increased expression of ß-Tubulin III at D60, indicative of successful neuronal differentiation. In comparison, while both the CCND2^E272*Het^ and CCND2^E272*Hom^ spheroids showed similar expression of ß-Tubulin III at D60, they also retained expression of PAX6 (Fig. 4), consistent with a defect in NP differentiation. CCND2^FS-T1^ and CCND2^FS-T2^ hCS also showed retained PAX6 expression at D60 but, in contrast to CCND2^E272^ hCS, showed little to no NeuN or ß-Tubulin III staining at D25 or 60.

**Figure 4.**
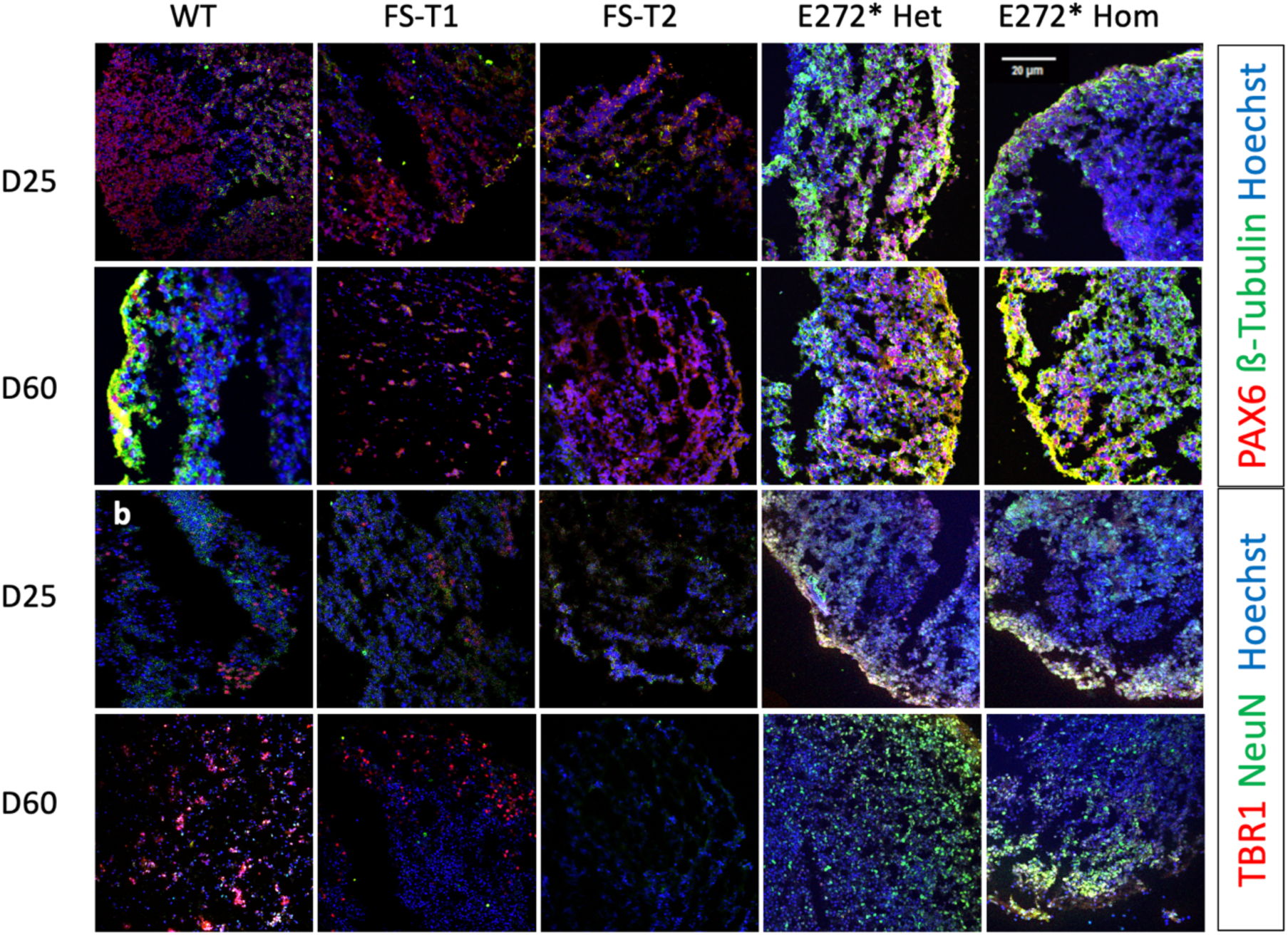
Representative fluorescence images of human cortical spheroids collected at D25 and D60 of differentiation following cryosectioning. Sections were stained for PAX6, β-Tubulin, TBR1, NeuN and Hoechst on all CCND2 CRISPR-Cas9 edited lines and compared to wildtype. For each, similar results were obtained from 2 independent batches grown. Scale bar 20um at x10 magnification.

IF staining using neuronal markers TBR-1 and NeuN revealed minimal staining across all hCS at D25 (Fig. 4). At D60 wildtype hCS showed positive staining for both NeuN and TBR-1, whereas CCND2^E272*Het^, CCND2^E272*Hom^ and CCND2^FS-T2^ hCS showed no evidence of TBR-1 expression, indicating a defect in neuronal differentiation and/or maturation. Interestingly, TBR-1 expression was observed in a proportion of cells in the D60 CCND2^FS-T1^ hCS. Overall, this indicates that CCND2 frameshift and nonsense mutations have different effects on neuronal expression in hCS.

In addition to changes in the expression of cell-type specific markers, we observed gross morphological differences between all the CCND2 edited lines compared to the wildtype, with the crispant lines generating spheroids that were less spherical in shape and developed “growth-pocket” like bulges by day 30 (Fig. 5).

**Figure 5.**
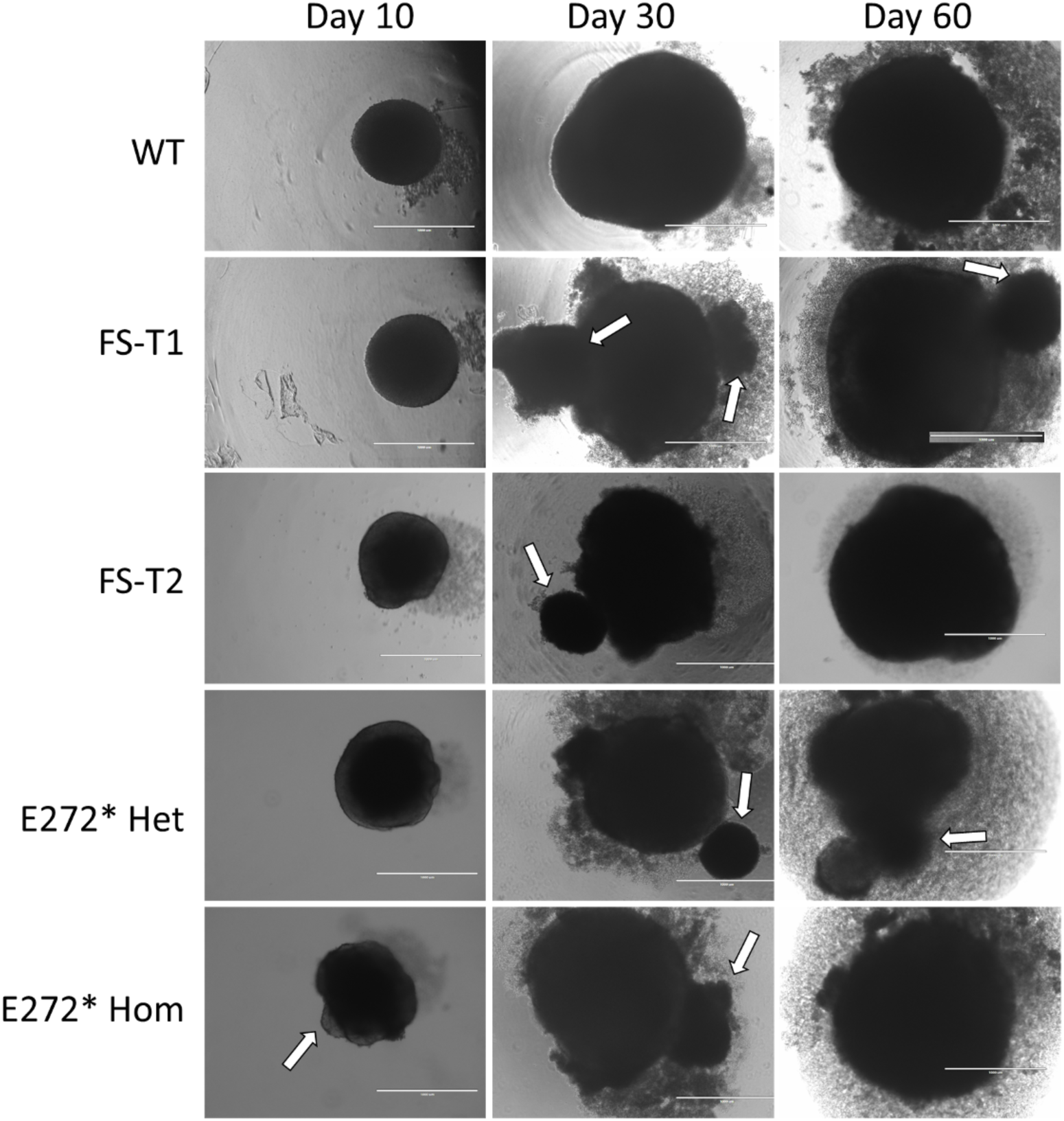
Human cortical spheroids with truncated CCND2 show morphological differences. A - Representative brightfield images of human cortical spheroids in culture at days 10, 30 and 60. For each line, similar results were obtained from 2 independent batches grown. Scale bar 1000um at x4 magnification. Arrows indicate “growth pocket” bulges.

## Discussion

Stabilisation of Cyclin D2 has been implicated in a range of human overgrowth disorders, specifically megalencephaly–associated syndromes such as MPPH and MCAP (Mirzaa et al., 2014, Zhao et al., 2024), as well as in haematological malignancies (Khanna et al., 2017). In order to investigate the mechanisms underpinning, and resulting from, CCND2 stabilisation during neurodevelopment, we generated four different isogenic human iPSC lines lacking the key regulatory Thr-280 residue at the c-terminus of CCND2. While we confirmed the MPPH-associated mutation, c.814G>T (p.Glu272*) resulted in CCND2 stabilisation in iPSCs, this was not the case for any of the CCND2 frameshift variants we generated.

Our findings are consistent with the literature, with no pathogenic frameshift CCND2 variants being reported as a cause of MPPH to date. Analysis of *CCND2* variants in ClinVar [accessed 01/05/2025] revealed 208 *CCND2* variants from which there are no frameshift variants. In comparison there are 8 nonsense variants, of which 7 are in the final exon, all located before Thr-280, and are considered pathogenic or likely pathogenic (p.Gln259Ter, p.Gln265Ter, p.Gly268Ter, p.Lys270Ter, p.Ser271Ter, p.Glu272Ter and p.Gln277Ter). Of note, frameshift and stop-gain mutations in the proximal region of *CCND2* have recently been reported to cause an inverse growth phenotype characterised by microcephaly, short stature, hypotonia and developmental delays (Pirozzi et al., 2021). Due to their location earlier in the gene, instead of stabilising CCND2 these *de novo* loss of function variants lead to a significant reduction in CCND2, likely triggering early cell cycle exit in neural progenitor cells.

Despite all the frameshift mutations removing the regulatory Thr-280 residue and impacting neurodevelopment in our hCS models, neither CCND2^FS-T1^ or CCND2^FS-T2^ accurately model MPPH due to their failure to cause CCND2 stabilization. This finding suggests that a lack of CCND2 Thr-280 is not in itself sufficient to cause CCND2 stabilisation. This is surprising given that single amino acid substitutions at Thr-280 and Pro-281 have both been shown to stabilise CCND2 (Mirzaa et al., 2014). One possible explanation is that the frameshift and premature termination codon negatively impact *CCND2* expression. The 3’ UTR of *CCND2* spans approximately 5,350bp and has been demonstrated to play a crucial role in regulating *CCND2* expression in the context of multiple myelomas. Similar to *CCND1*, the 3’UTR of *CCND2* contains binding sequences for over 255 miRNA, with reductions in UTR size correlating with increased *CCND2* expression due to loss of miRNA binding sites (Misiewicz-Krzeminska et al., 2016). The premature truncations caused by the frameshift mutations we introduced may therefore result in an earlier start position of the 3’ UTR, directly after the PTC, resulting in an increase in the length of the 3’UTR, corresponding with reduced *CCND2* expression. Care should therefore be taken when assessing and interpreting variants based on functional assays using exogenous *CCND2*, which lack both introns and UTRs, as these may not accurately reflect CCND2 regulation and stabilisation.

Human cortical spheroids mirror foetal neuronal development and mimic similar megalencephaly-associated overgrowth disorders, such as those caused by mutation of TSC1/2 (Blair et al., 2018). Mouse models have similarly modelled brain overgrowth conditions, showing that mutations in the PI3K-AKT-mTOR pathway accurately recapitulate phenotypes seen in patients such as megalencephaly, cortical migration defects and medically refractive epilepsy (Roy et al., 2015, Groszer et al., 2001). The megalencephaly phenotype, in particular, results from an increased number of proliferating PAX6^+^ neural progenitor cells (Li et al., 2017) which reside in the ventricular and subventricular zones of the developing cortex (Kikkawa et al., 2019). In D25 hCS, we observed few differences between wildtype and crispant lines, with all mutants displaying similar numbers of PAX6 positive cells compared to the wildtype. However, at D60, both the frameshift and nonsense hCS had significantly more PAX6^+^ cells compared to the wildtype, indicating a larger proportion of NP cells remained in a proliferative state and had failed to differentiate. Furthermore, the few remaining PAX6 positive cells in the wildtype hCS had migrated to the “cortical plate” edge of the spheroid along with the β Tubulin III positive neurons, whereas in CCND2 crispant spheroids the PAX6 positive cells were more densely spread throughout mimicking similar findings to other macrocephaly associated disease models (Li et al., 2017). PAX6 plays a multifunctional role in neurogenesis, determining neural progenitor fate as well as proliferation (Zhang and Jiao, 2015). PAX6 expression is paused in early postnatal periods of healthy embryos (Brill et al., 2009), followed by the subsequent reduction of PAX6 expression as NP cells terminally differentiate in the mature brain (Ninkovic et al., 2010). The persistence and increased number of PAX6 expressing cells, is indicative of uncontrolled neural progenitor cell proliferation, and is seen in organoid and mouse models of MCAP and MPPH (Roy et al., 2015, Zhang et al., 2020). The finding of persistent PAX6 expression in our CCND2 hCS is therefore in keeping with existing models and fits with the ongoing hypothesis that progenitor proliferation defects are seen in patients with megalencephaly associated disorders (Guarnieri et al., 2018).

Also involved in neuronal migration, the transcription factor TBR-1 regulates early born neurons during embryogenesis (Zhang and Jiao, 2015, Han et al., 2011) and is not present in neural progenitor cells (Hevner et al., 2001). As expected at D60, TBR-1 expression was observed in wildtype hCS, which should reach peak expression by day 76 (Pasca et al., 2015). However, no TBR-1 expression was observed in either CCND2^E272*Het^ or CCND2^E272*Hom^ hCS. Despite an earlier truncation at position 266 in the frameshift line CCND2^FS-T1^ compared to position 270 in CCND2^FS-T2^, a complete absence of TBR-1 positive cells was observed in CCND2^FS-T2^ hCS, whereas TBR-1 positive cells were present in CCND2^FS-T1^ hCS. Despite the close proximity in the position of the termination codon between the two frameshift mutant lines, CCND2^FS-T1^ and CCND2^FS-T2^ appeared to model neurogenesis differently, with a stronger impact observed in CCND2^FS-T2^. However, neither frameshift nor nonsense mutant hCS were consistent with the wildtype differentiation at D60.

In addition to Cyclin D2 not being stabilized in CCND2^FS-T1^ and CCND2^FS-T2^, both frameshift lines showed very little neuronal maturation with minimal expression of ß-Tubulin III and NeuN compared to the wildtype. In comparison, both CCND2^E272*Het^ and CCND2^E272*Hom^ lines displayed Cyclin D2 stabilization and an increased pool of PAX6 positive neural progenitor cells which failed to differentiate into mature TBR-1 positive neurons in D60 hCS, all features consistent with MPPH. Despite the close proximity of the truncations, our findings indicate subtle differences in how CCND2 frameshift and nonsense mutations impact neurogenesis, with nonsense mutations better mimicking MPPH.

In summary, we have created four crispant CCND2 truncating iPSC lines and differentiated them into human cortical spheroids to model MPPH-associated neurodevelopmental disease. Our data confirms that the heterozygous CCND2^E272*^ line is a representative model of MPPH. While the homozygous CCND2^E272*^ line did mimic the phenotypes observed in MPPH, to date no patients with biallelic pathogenic *CCND2* variants have been reported. The lack of CCND2 stabilisation in the frameshift lines provides insight as to why no MPPH patients have been identified with a frameshift truncation in the final exon of CCND2, despite them resulting in a similar truncated protein to nonsense mutations, lacking the Thr-280 residue. Further work to characterise these models is required to understand the role of CCND2 dysregulation in MPPH and associated neurodevelopmental conditions to identify possible treatment options.

## Materials and Methods

### Culture of iPSCs

AD2 iPSCs were a generous gift from Professor Majlinda Lako. iPSCs were cultured in complete mTeSR plus media (Stem Cell Technologies) on Matrigel® coated (Corning) ultra-low attachment 6 well plates (Corning) and passaged with ReLeSR (Stem Cell Technologies) at approximately 70% confluency. Cultures were regularly tested for pluripotency by Immunofluoresence of Oct3/4 and SSEA, and mycoplasma using MycoStrips (Invitrogen).

### CRISPR-Cas9 genome editing of iPSCs

A single-guide RNA targeting the final exon of CCND2 (5’-ACGTGACGGATCCAAGTCGG-3’) was ligated in to either the GFP-tagged CRISPR-Cas9 px458 plasmid (Addgene #48138) or the enhanced specificity Cas9 (eSpCas9, Addgene #71814) following digestion with BbsI restriction enzyme (Thermo Fisher Scientific). Sanger sequencing was performed to confirm the successful ligation of the guide. A single-stranded oligodeoxynucleotide (ssODN) repair template designed to introduce the specific nonsense mutation CCND2 E272* was manufactured by Integrated DNA Technologies. (5’CTGCAGCAGTACCGTCAGGACCAACGTGACGGATCCAAATCCGAGGATGAACTGGACCAAGCCAGCGCCCCTACAGACGTGCGGGATATCGACCTGTGAGGATGCCA-3’).

For each reaction, 2µg of sequence validated Cas9 plasmid, and 2-6 µg of ssODN repair template, when required, was mixed with 800,000 wildtype iPSCs lifted with 0.75X TrypleE (Thermo Fisher Scientific) and electroporated using the Human Stem Cell Nucleofector^TM^ Kit (Lonza). After electroporation, solutions were transferred to 24-well plates containing mTESR plus and 10uM ROCK inhibitor Y-27632 (Stratech Scientific). 48 hours later, cells were detached using 0.75X TryplE and resuspended with agitation in mTeSR plus containing 0.1% BSA and 10uM ROCK inhibitor Y-27632, before using a 40µm filter to create a single cell suspension. The resulting suspension was FACS sorted for GFP positive cells into Matrigel coated 96-well plates, with 1 GFP-positive cell per well. Cells were incubated for 48hrs in mTeSR supplemented with 10µM of ROCK inhibitor Y-27632 and 1% penicillin/streptomycin, followed by a media change to mTESR plus alone. Successful colonies were propagated for 7-10 days and genomic DNA extracted using DirectPCR Lysis Reagent (Viagen). PCR amplification of CCND2 exon 5 was performed (Forward primer: 5’ TGCTCTATGTCCTGTTCCTCT 3’ and Reverse primer: 5’ TGTTCGCATATACAAGGGATTCC 3’) and Sanger sequenced to identify colonies harbouring mutations. TOPO cloning (Thermo Fisher Scientific) was performed for the colonies with biallelic frameshift mutations to confirm the exact sequence of each allele.

### Immunocytochemistry of iPSC pluripotency makers

iPSCs were seeded on Matrigel®-coated glass coverslips and cultured until 60% confluent. Cells were subsequently fixed in 4% paraformaldehyde (PFA) and permeabilised in 0.2% Triton-X-100 [v/v] in PBS. Following blocking with 3% bovine serum albumin in PBS, primary antibodies were incubated for 2 hours at their optimal dilution in 3% bovine serum albumin. Coverslips were washed 3 times with 0.05% Triton in PBS, followed by a final 1x PBS-only wash and then incubated with fluorescently labelled AlexaFluor secondary antibodies (1/2,000) and Hoechst (Thermo Fisher Scientific) for 1hr at room temperature. Three final washes with 0.05% Triton in PBS, followed by a final 1x PBS-only wash were performed before coverslips were mounted on microscope slides using Fluoromount mounting medium (Thermo Fisher Scientific). The following antibodies were used for assessment of pluripotency; Goat-Oct 3/4 (1/30, R&D Systems; AF1759), Alexa Fluor Dye donkey anti-goat-IgG AlexaFluor488 (1/500) and Alexa Fluor® 555 Mouse anti-SSEA-4 (1/50, BD Biosciences; 560218). Once cured, cells were imaged on a Nikon A1R confocal microscope and the resulting images processed using ImageJ software.

### Cycloheximide assay in cultured iPSCs

To assess Cyclin D2 stabilisation, iPSCs were cultured in mTESR plus media containing 10ug/ml cycloheximide for 90 minutes. This was done on each iPSC line at 60% confluency while cells were actively proliferating. Cells were subsequently scraped and lysed using NP-40 lysis buffer containing 1 x Halt protease and phosphatase inhibitors (Thermo Fisher Scientific). Protein lysates were quantified using the Biorad DC kit (BioRad) then Cyclin D2 stabilisation assessed by western blotting.

### Western blotting

Cultured iPSCs were detached with ReLeSR and resuspended in NP-40 lysis buffer containing 1X Halt protease and phosphatase inhibitors. Protein concentration was determined using the BioRad DC kit. 30μg of protein in 4x LDS sample buffer (Thermo Fisher Scientific) was loaded onto a 4-12% Bis-Tris Gel (Thermo Fisher Scientific) and transferred to polyvinylidene difluoride membranes following methanol activation. Membranes were blocked in 5% skimmed milk for 2 hours at room temperature and incubated overnight at 4°C with primary antibodies diluted in skimmed 1% milk. The following day, membranes were washed in 1X PBS and incubated with HRP-conjugated secondary antibodies for 1 hour at room temperature.: Rabbit-CCND2 (CST, 1/1,000) and Mouse-B-actin (Ambion, 1/20,000) antibodies were used. Following three washes in 1X PBS, bound antibodies were viewed using SuperSignal West Femto Chemiluminescent substrate on a Bio-Rad Chemi-Doc imaging system with a chemiluminescent filter. Images were taken, and densitometry analysis performed using ImageLab software (BioRad). Where required, following imaging, blots were stripped using Restore^TM^ Western Blot Stripping Buffer (Fisher Scientific) and re-blocked and blotted with additional antibodies. Secondary antibodies used were Goat Anti-Rabbit HRP (#7074) or Horse anti-mouse HRP (#7076) at a final dilution of 1/5,000.

### Differentiation and culture of 3D cortical spheroids

Differentiation of iPSCs into 3D cortical spheroids was performed as described previously (Pasca et al., 2015). Briefly, confluent undifferentiated iPSC colonies were detached with Accutase and resuspended in complete TeSR-E6 media (Stem Cell Technologies), supplemented with 10µM of ROCK inhibitor Y-27632. 15,000 cells were seeded per well of a 96-well, Lipidure coated low-adhesion U bottom plate (AMSBIO) and incubated for 24 hours (day 0) in an incubator at 37°C and 5% CO2. On days 1-5, the media was changed daily to complete TeSR-E6 media, supplemented with 2.5µM Dorsomorphin (also known as compound C; Sigma) and 10µM SB431542 (Generon). On day 6, the media was changed to neural induction media composed of Neurocult (Stem Cell Technologies), B-27 Supplement without vitamin A (Thermo Fisher Scientific), 100U/ml penicillin-streptomycin, and 1X GlutaMAX (Thermo Fisher Scientific), supplemented with 20ng/ml FGF (R&D Systems) and 20ng/ml EGF (R&D Systems). Spheroids were fed with neural induction media for 19 days, with daily medium changes for the first 10 days, and every other data for the following 9 days. From days 25-43, the developing spheroids were grown in neural induction media supplemented with 20ng/ml BDNF (Peprotech) and 20ng/ml NT-3 (Peprotech), with media changes every 2 days. From day 43, spheroids were maintained in neural induction media without supplements, with media changes every 4 days until harvest.

### Immunofluorescent analysis of cortical spheroids

Cortical spheroids were removed from growth media and washed in 1x PBS before being fixed in 4% PFA at 4°C overnight. Following fixation, spheroids were washed in PBS and dehydrated for 72 hours at 4°C in 30% sucrose before being embedded in OCT compound (Agar Scientific), snap frozen on dry ice and stored at -80°C. For immuno-staining, 8µM sections were cut from the resulting tissue blocks on a cryostat and washed once with PBS to remove excess OCT. For immunocytochemistry, sections were hydrated in PBS and permeabilised with 0.2% Triton-X-100 [v/v] in PBS. Sections were washed again in PBS and then blocked in 3% bovine serum albumin for 1 hour at room temperature. Sections were incubated in primary antibodies diluted in 3% bovine serum albumin overnight at 4°C. Primary antibodies used were: Rabbit PAX6 (BioLegend, #901301, 1/100), Mouse β-Tubulin III (Sigma, #T8578, 1/100), Rabbit Tbr-1 (Millipore, #AB10554, 1/100) and Mouse NeuN (Millipore, #MAB377, 1/30). The following day, sections were washed three times with 0.05% Triton in PBS, followed by a single 1x PBS wash and incubated with secondary antibodies (1/500, Alexa Fluor Dyes (donkey anti-rabbit-IgG AlexaFlour488; Donkey anti-mouse-IgG AlexaFluor568) (Thermo Fisher Scientific) and Hoechst (1/200) diluted in 3% bovine serum albumin for 1 hour at room temperature. After repeated washes in 0.05% Triton in PBS and 1x PBS, slides were cover-slipped using Fluoromount mounting medium and allowed to set overnight before imaging. Images were taken on a Nikon A1R confocal microscope, using 10x, 20x or 40x objectives and processed using ImageJ software.

## Acknowledgements

pSpCas9(BB)-2A-GFP)PX458) was a gift from Feng Zhang (Addgene plasmid #48138 (Ran et al., 2013)). eSpCas9(1.1) was a gift from Feng Zhang (Addgene plasmid # 71814 (Slaymaker et al., 2016).

## Data Availability

The iPSC lines generated in this study are available on request from the corresponding author.

## Author contributions

E.L.H and J.A.P. conceived the idea of the study. E.L.H., R.D.T., A.M.S.R and K.S. conducted laboratory experiments. J.A.P and E.G.S. obtained funding for the study. K.S., J.A.B, C.A.J. and J.A.P. supervised the research. E.L.H. and J.A.P. wrote the manuscript draft. All authors read and approved the manuscript prior to submission.

